# OBO Foundry in 2021: Operationalizing Open Data Principles to Evaluate Ontologies

**DOI:** 10.1101/2021.06.01.446587

**Authors:** Rebecca C. Jackson, Nicolas Matentzoglu, James A. Overton, Randi Vita, James P. Balhoff, Pier Luigi Buttigieg, Seth Carbon, Melanie Courtot, Alexander D. Diehl, Damion Dooley, William Duncan, Nomi L. Harris, Melissa A. Haendel, Suzanna E. Lewis, Darren A. Natale, David Osumi-Sutherland, Alan Ruttenberg, Lynn M. Schriml, Barry Smith, Christian J. Stoeckert, Nicole A. Vasilevsky, Ramona L. Walls, Jie Zheng, Christopher J. Mungall, Bjoern Peters

## Abstract

Biological ontologies are used to organize, curate, and interpret the vast quantities of data arising from biological experiments. While this works well when using a single ontology, integrating multiple ontologies can be problematic, as they are developed independently, which can lead to incompatibilities. The Open Biological and Biomedical Ontologies (OBO) Foundry was created to address this by facilitating the development, harmonization, application, and sharing of ontologies, guided by a set of overarching principles. One challenge in reaching these goals was that the OBO principles were not originally encoded in a precise fashion, and interpretation was subjective. Here we show how we have addressed this by formally encoding the OBO principles as operational rules and implementing a suite of automated validation checks and a dashboard for objectively evaluating each ontology’s compliance with each principle. This entailed a substantial effort to curate metadata across all ontologies and to coordinate with individual stakeholders. We have applied these checks across the full OBO suite of ontologies, revealing areas where individual ontologies require changes to conform to our principles. Our work demonstrates how a sizable federated community can be organized and evaluated on objective criteria that help improve overall quality and interoperability, which is vital for the sustenance of the OBO project and towards the overall goals of making data FAIR.

## Introduction

The quantity and complexity of data generated by biological experiments are growing at an unprecedented rate. Ontologies are used to organize, annotate, and analyze this data, and to harmonize the rich and varied information captured in key biological knowledge bases (1). A major challenge faced by researchers is the large numbers of different overlapping ontologies, varying in quality and completeness, each attempting to cover different aspects of any given domain of interest. For example, BioPortal (2) includes over 800 ontologies and close to ten million terms as of April 2021 (https://bioportal.bioontology.org/). These challenges are compounded when we consider the fact that many applications require using *combinations* of ontologies. If ontologies are constructed using different principles, they will not work together in a modular, interoperable, coherent way.

The Open Biological and Biomedical Ontologies (OBO) project was initiated in the early 2000s, as it became clear that there was a community desire to expand ontologies beyond the scope of the Gene Ontology (GO) to tackle biological and biomedical problems more broadly (3). OBO was designed to organize and guide the development of ontologies according to common standards and principles (4), enabling modular composition of ontologies and providing guarantees of technical and scientific quality. One of the mechanisms was a set of **principles**, which were to be followed by all ontologies within the OBO Foundry (**Figure 1**). For example, OBO ontologies should be *open*, allowing for reuse, and the ontologies should conform to shared standards for how terms are interrelated. Currently, OBO is governed by a volunteer team consisting of ontology maintainers and stakeholders (the ‘OBO operations committee’), represented by the authors of this manuscript. This team carries out multiple duties, including maintaining the site, stewarding the principles, and curating ontology metadata.

Here we describe our efforts to **operationalize the OBO Foundry principles**. Working closely with stakeholders across OBO, we have refined the original principles, codifying them into operational tests that can be executed automatically at regular intervals. We have implemented a dashboard that provides a matrix view indicating the conformance to each principle for each of the over 150 active ontologies in OBO, allowing drill-down to see complete reports. This work involved significant community effort, working with individual ontologies, and required a wholesale re-curation of ontology metadata across OBO. The results allow both ontology developers and the broader community of users to see the steps each ontology must take to come into conformance.

## Results

### Capturing consistent ontology metadata in the OBO registry

OBO considers two sources of information for each ontology project: The ontology itself and metadata provided by the ontology maintainers stored in the OBO registry (http://obofoundry.org/). To automate the evaluation of principles across OBO ontologies, we first wanted to ensure that the OBO registry entries accurately and consistently captured the minimal information listed in **Table 1**. The OBO registry has grown from a short and simple list of a dozen ontologies to a comprehensive resource for metadata on more than 150 active projects. To ensure that the information in the OBO registry was up to date, we emailed the indicated contact persons for each ontology. If no response was obtained, we used personal contacts as well as searches on PubMed and Google to try to find alternative contacts. Overall, we found that out of 201 ontologies, 145 were under current active development, 5 were in use but not being actively developed, 45 were obsoleted, and for 6, no contact person could be identified, making them ‘orphaned’. For the active ontologies, we asked the developers to confirm and update fields in the OBO registry, specifically the ontology title, homepage, contact, description, and license. This resulted in a total of over 60 updates to OBO registry metadata, most of which were additions of previously missing information.

To ensure that the OBO registry records will be kept up to date over time, we created a lightweight system for collaboratively curating and updating these records. Metadata files are stored in a structured format under version control in a repository within the OBO GitHub organization. This allows both ontology maintainers and members of the core OBO team to make suggestions via GitHub pull requests. This metadata is visible to the community via the OBO registry website, or in computable format (YAML and JSON-LD), and is used in order to evaluate an ontology according to the newly operationalized principles. As of May 2021, there have been 3045 commits by 113 developers to the new repository, demonstrating that this system is adequate for broad use by the OBO community. The end result of this process is consistent and quality-controlled metadata for each ontology, and a procedure for ensuring these can be easily kept up-to-date by the community.

### Defining operating principles for OBO ontologies

We took the original set of OBO principles and for each one, refined them until we had arrived at a more crisply stated operational procedure. For example, the first principle of OBO is that the ontology is ‘open’. However, there were no specific recommendations on the licensing terms that would meet that goal, or of how the license should be stated. Some ontologies included license information on their home page, others embedded it in their ontology metadata. After community discussions, we agreed that ontologies could be considered ‘open’ for the purposes of OBO if they used the Creative Commons Attribution (CC BY) license 3.0 or later, or if they were in the public domain using the Creative Commons CC0 declaration. Both of these options conform to the spirit of the original principle of openness, and were already adopted widely by a majority of OBO ontologies. Next, we settled on a convention on how the license should be stated, and decided on the use of the widely accepted Dublin Core Terms (5) ‘license’ property (“dcterms:license”) in the ontology file metadata in addition to a declaration of the license in the OBO registry entry. These conventions allow checking for the presence of an ‘Open’ license computationally, in both the ontology file itself and the information contained in the OBO registry.

Following the same process for each principle, **Table 2** lists how each principle is now encoded with a succinct summary of the principle using ISO MUST/SHOULD language (6) (https://tools.ietf.org/html/rfc2119), and a description of the automated check being performed. A more detailed description of each principle is linked to, which includes a description of the Purpose (what the principle is intended to achieve); *Recommendations* for ontology developers describing how they should best conform to the principle; examples of *Implementation* of the principle; *Counter examples* showing how an ontology could fall short of conformance to the principle; and *Criteria for review* that spell out what a human reviewer should be looking for in an ontology in order to judge if it adheres to the principle or not. Each principle has a corresponding issue on the public GitHub repository (https://github.com/OBOFoundry/OBOFoundry.github.io) in which further questions and discussions are tracked, and there is a continuous review in bi-weekly conference calls of new questions and the need to update the wording of principles. At the same time, anyone is able to asynchronously comment on the process by adding their comments to the relevant GitHub issue.

### Establishing a framework for automatic evaluation of ontologies

In order to semi-automate the process of determining ontology conformance, we implemented a validation suite that displays its results through the OBO dashboard (http://dashboard.obofoundry.org/dashboard/index.html). The dashboard implements an executable programmatic expression of each principle, and a framework for running these checks, and for delivering a web-based report. The dashboard is implemented on top of the ROBOT software suite (7), and in particular, uses the ability of ROBOT to reason over ontologies and to generate detailed reports. Additionally, the validation suite checks the metadata for each ontology in the OBO registry. For example, the curated ‘usages’ tag is used to determine if the ontology fulfills the criterion for having a plurality of independent users.

The dashboard results are shown as a grid where each ontology is a row and each OBO principle a column, with each cell indicating results of the check for this combination (**Figure 2**). For each OBO principle, the dashboard links to 1) the web page for that principle, which links to 2) a web page describing the automated test, which links to 3) a tracker issue for the automated test. Each ontology has a detailed report page accessible from the main dashboard by clicking on the ontology ID. This provides a breakdown of the problems encountered and suggestions on how to fix them.

When a preliminary version of the dashboard was first announced to the OBO ontology maintainers in early 2020, several ontology maintainers started fixing the problems identified in the dashboard scripts. Specifically, comparing the experimental dashboard runs in 11/2019 (prior to the announcement of the OBO dashboard work) vs. 07/2020, we found a significant reduction in reported errors identified by the dashboard code (p=0.0005, Wilcoxon matched pairs test, **Figure 3**). At the same time, users reported issues with the automated validation code leading to false-positive and false-negative results, which were subsequently fixed and have led to the more robust version of the code implemented in the current version of the dashboard. While the iterative updates to the code mean that current numbers of validation issues cannot be compared to those at the start of the project, the community engagement and the noticeable drop in issues between versions that could be compared demonstrate that the OBO ontology developer community is responsive to the issues identified by the dashboard, and that highlighting problems in a transparent manner can be a productive first step to resolving them.

As can be seen in **Figure 4**, as of May 2021, four principles are fully conformed to by all 175 OBO Foundry ontologies: *FP02 Common Format, FP03 URIs, FP11 Locus of Authority*, and *FP20 Responsiveness*. The principle that is least conformed to is *FP06 Textual Definitions*, with only 19 ontologies (about 11%) fully passing this check.

## Discussion

The scientific community has always relied on sharing data through publications or personal communications. The recently developed FAIR principles (8) spell out what it takes for shared data to be Findable, Accessible, Interoperable, and Reproducible. A key requirement of FAIR is to use vocabularies that are reusable across projects, which aligns with the original goals of the OBO project, which precedes the formulation of the FAIR principles by more than a decade. Thus, the goals of OBO and FAIR are highly compatible, and the lessons learned from our work on OBO should be taken into consideration when evaluating FAIR principles.

Like FAIR, the original OBO principles served as a rallying cry, galvanizing a community to work towards a broadly articulated vision. After two decades of work on OBO, we found that relying on human review of such principles is difficult to standardize and does not scale. Instead, we decided to turn each principle into operational tests for conformance. We found that this process was beneficial to communicating clearly what each principle was meant to accomplish and to provide clear guidance for ontology developers on what they needed to do to achieve compliance with the principle.

Going forward, we plan to run the OBO Dashboard on all new ontologies requesting OBO membership, and on each new release of every OBO member project. Given the free availability of the code, it can be run (and in some cases already is running) as part of internal ontology development pipelines to test internal release candidates. We expect that this process will identify weaknesses in the current pipeline, and result in continuous improvements of the tests themselves, and of the shared understanding of what the tests (and the principles) are meant to achieve across the OBO community.

There are several limitations to our approach. First, the current framework examines a single ontology at a time. We are planning to extend the checks to run across sets of ontologies to provide insights on inter-ontology consistency. Second, not all principles formulated for the OBO Foundry can be checked reliably in an automated fashion. Specifically, human review is needed to check for scope, a plurality of users, and co-operation with existing ontologies. While these limitations have to be kept in mind, it is important to realize how much more consistent and up-to-date the current automated system is compared to the previous practice of relying on manual human volunteer reviewers.

In conclusion, this manuscript highlights the OBO dashboard and associated automated test as the main advancement of the OBO Foundry in 2021. As this is the first official publication of the OBO dashboard, we expect that there will be community feedback and criticism on the specific implementation of the checks implemented, and we very much welcome that. We hope that the quantitative nature of the dashboard and its underlying automated rules will make these discussions constructive. Furthermore, we hope that other standardization-focused projects will take inspiration from the OBO Foundry’s successful effort to assess and quantify our evaluation principles.

## Acknowledgments

This work was supported by NIH grant R24HG010032. We thank the members of the OBO community who have provided feedback on the dashboard, including Chris Grove, Ceri van Slyke, Sebastian Koehler, James Seager, Anne Thessen, Sue Bello, Midori Harris, Hilmar Lapp, Charles T Hoyt. The work of CM, WD, SC, SL, and NH was supported in part by the Director, Office of Science, Office of Basic Energy Sciences, of the U.S. Department of Energy under Contract No. DE-AC02-05CH11231.

## References

1. Blake JA and Bult CJ. (2006) Beyond the data deluge: data integration and bio-ontologies. J. Biomed Inform., 39(3), 314–320.

2. Whetzel PL, Noy NF, Shah NH, et al. (2011) BioPortal: enhanced functionality via new Web services from the National Center for Biomedical Ontology to access and use ontologies in software applications. Nucleic Acids Res., 39(Web Server issue), W541–5.

3. Ashburner M, Mungall CJ, and Lewis SE. (2003) Ontologies for biologists: a community model for the annotation of genomic data. Cold Spring Harb Symp Quant Biol., 68, 227–235.

4. Smith B, Ashburner M, Rosse C, et al. (2007) The OBO Foundry: coordinated evolution of ontologies to support biomedical data integration. Nat Biotechnol., 25(7), 1251–1255.

5. Weibel S, Kunze J, Lagoze C, and Wolf M. (1998) Dublin core metadata for resource discovery. Internet Engineering Task Force RFC, 2413(222), 132.

6. Bradner S. (1997) RFC2119: Key words for use in RFCs to Indicate Requirement Levels. USA: RFC Editor, 10.17487/RFC2119.

7. Jackson RC, Balhoff JP, Douglass E, et al. (2019) ROBOT: A Tool for Automating Ontology Workflows. BMC Bioinformatics, 20(1), 407.

8. Wilkinson MD, Dumontier M, Aalbersberg IJJ, et al. (2016) The FAIR Guiding Principles for scientific data management and stewardship. Sci. Data, 3, 160018.

