## Supplementary material for "OBO Foundry in 2021: Operationalizing Open Data Principles to Evaluate Ontologies": Figures

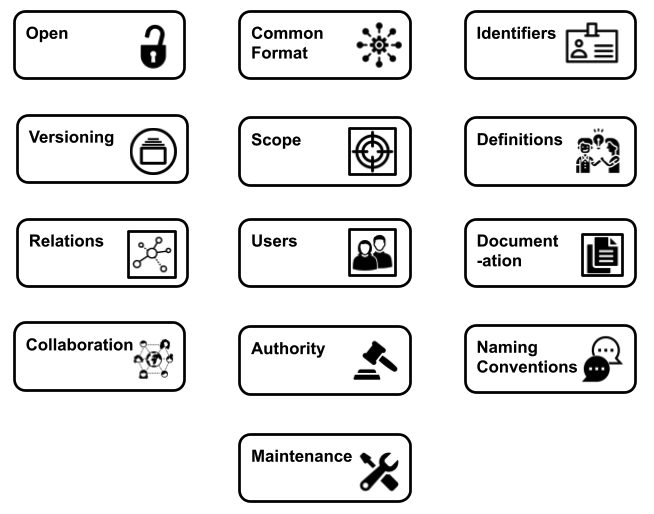

**Figure 1:** Illustration of the principles around which the OBO Foundry was built

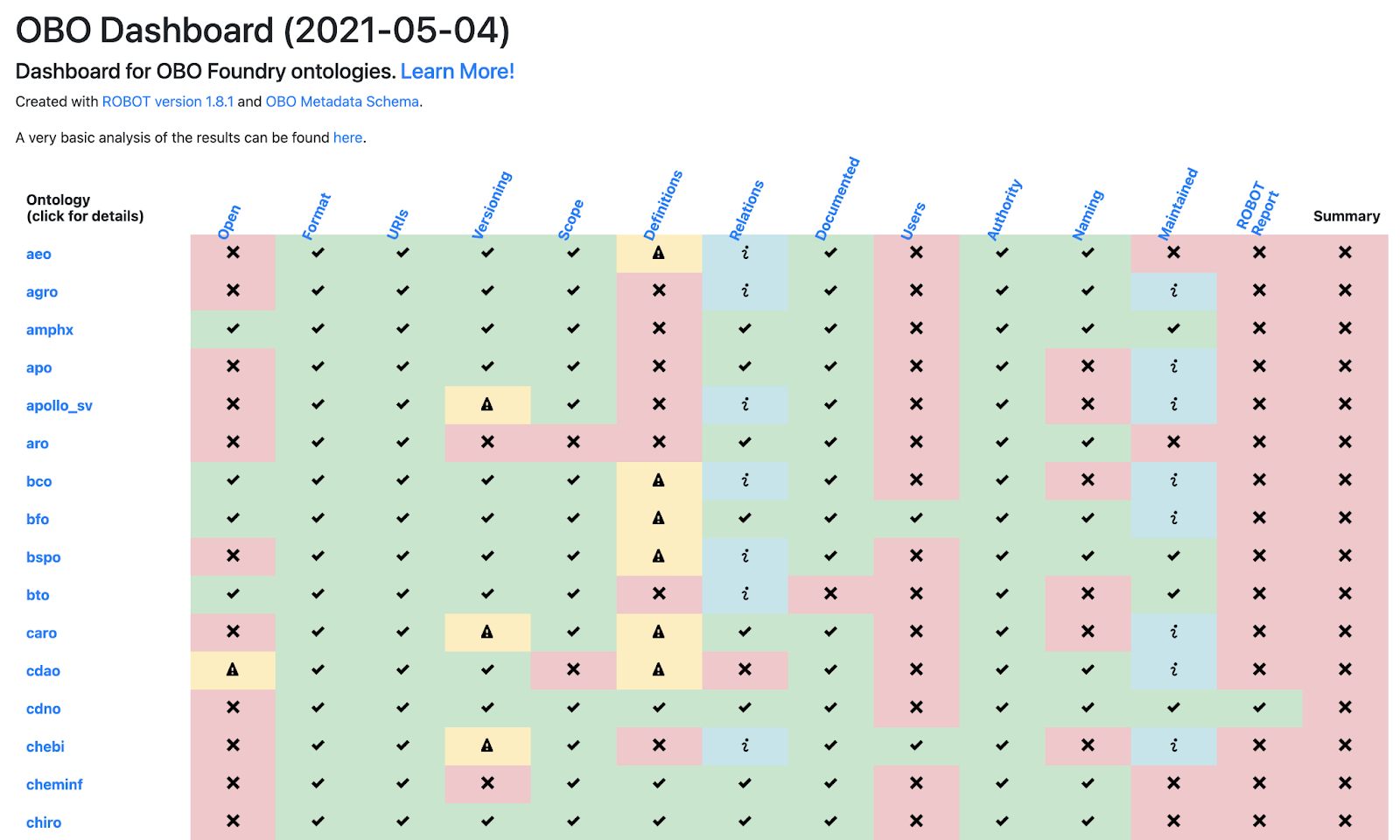

**Figure 2**: The OBO Dashboard (truncated). The rows represent OBO ontologies (of which the first 15 in alphabetical order are shown here) and the columns are the OBO principles. The final column (Summary) shows whether the ontology passed all of the tests. Clicking on the ontology ID in the far left column directs to a detailed report page.

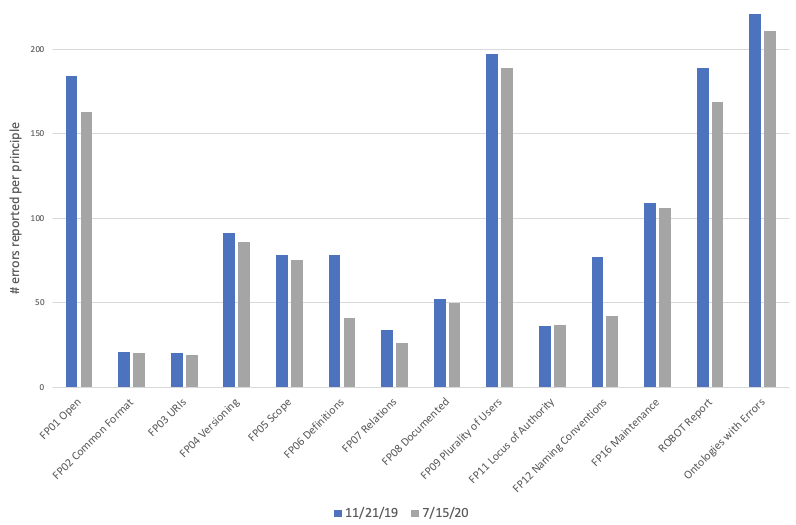

**Figure 3:** Number of errors reported by dashboard in November 11, 2019 (blue bars) and July 15, 2020 (gray bars). The final column, “Ontologies with Errors”, is the total number of ontologies that had one or more errors, not a count of all errors.

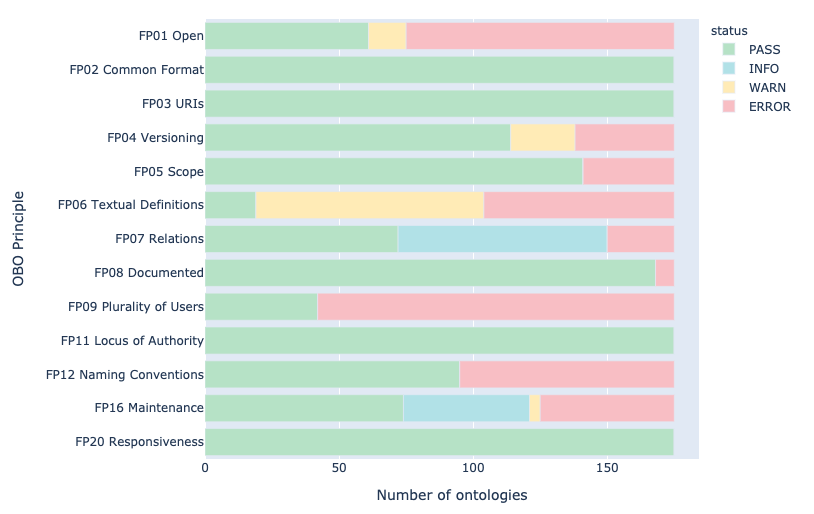

**Figure 4:** Summary of principle conformance across all OBO Foundry ontologies in May 2021
