## Supplementary material for "OBO Foundry in 2021: Operationalizing Open Data Principles to Evaluate Ontologies": Tables

| **Field** | **Definition** | **Example** | **Automated validation** |
| --- | --- | --- | --- |
| title | Full name of the ontology | Ontology for Biomedical Investigations | must be present |
| id | Abbreviation of the ontology’s name used as the exclusive namespace for the ontology | obi | lowercase, alphanumeric, no spaces |
| homepage | Website where a user can find information about the ontology | <http://obi-ontology.org/> | must be in URL format |
| contact.label | Name of a person responsible for the ontology | Bjoern Peters | must not contain ‘@’ and only have one label |
| contact.email | Email address of the person responsible for the ontology | | must be in email format and only have one email |
| products.id | Name of the canonical ontology file (id.owl). Additional products may include ontology subsets, bridge files, etc. | obi.owl | format enforced |
| description | Concise free text description of the scope of the ontology | An integrated ontology for the description of life-science and clinical investigations | must be present |
| license.label | Name of license | CC-BY 4.0 | must correspond to the title of the license.url |
| license.url | URL of license | https://creativecommons.org/licenses/by/4.0/ | must be in URL format |
| activity status | Indicates the development status of the ontology | active | must be one of ‘active’, ‘inactive’, or ‘orphaned’ |
| obsoletion status | Indicates if the ontology has been declared obsolete by the original developers | false | must be either true or false |

**Table 1: Minimal ontology metadata captured in the OBO registry**

| **Principle** | **Summary** | **Automated Check** |
| --- | --- | --- |
| Open (<http://obofoundry.org/principles/fp-001-open.html>) | The ontology MUST be openly available to be used by all without any constraint other than (a) its origin must be acknowledged and (b) it is not to be altered and subsequently redistributed in altered form under the original name or with the same identifiers. | The registry data entry is validated with JSON Schema. The license schema ensures that a license entry is present and that the entry has a url and label. The schema also checks that the license is one of the CC0 or CC-BY licenses. Then, annotations from the ontology are retrieved and the “dcterms:license” annotation is retrieved (if exists). The script ensures that the correct “dcterms:license” property is used and compares this license to the registry license to ensure that they are the same. |
| Common Format (<http://obofoundry.org/principles/fp-002-format.html>) | The ontology is made available in a common formal language in an accepted concrete syntax. | The ontology is loaded using OWLAPI. If the ontology is successfully loaded, it is assumed that it is in a good format. |
| URI/Identifier Space (<http://obofoundry.org/principles/fp-003-uris.html>) | Each class and relation (property) in the ontology must have a unique URI identifier that follows the format: a base URI + a prefix that is unique within the Foundry + a local numeric identifier. | All entity IRIs are retrieved from the ontology, excluding annotation properties. Annotation properties may use hashtags and words due to legacy OBO conversions for subset properties. All other IRIs are checked if they are in the ontology’s namespace. If the IRI begins with the ontology namespace, the next character must be an underscore. The IRI is also compared to a regex pattern to check if the local ID after the underscore is numeric. |
| Versioning (<http://obofoundry.org/principles/fp-004-versioning.html>) | The ontology provider has documented procedures for versioning the ontology, and different versions of ontology are marked, stored, and officially released. | The version IRI is retrieved from the ontology and, if found, this IRI is compared to a regex pattern to determine if it is in date format. |
| Scope (<http://obofoundry.org/principles/fp-005-delineated-content.html>) | The scope of an ontology is the extent of the domain or subject matter it intends to cover. The ontology must have a clearly specified scope and content that adheres to that scope. | First, the registry data is checked for a ‘domain’ tag. If it is present, the domain is compared to all other ontology domains. If the ontology shares a domain with one or more other ontologies, we return a list of those ontologies. |
| Textual Definitions (<http://obofoundry.org/principles/fp-006-textual-definitions.html>) | The ontology has textual definitions for the majority of its classes and for top level terms in particular. | ROBOT “report” is run over the ontology. A count of violations for each of the following checks is retrieved from the report object: duplicate definition, multiple definitions, and missing definition. |
| Relations (<http://obofoundry.org/principles/fp-007-relations.html>) | Relations should be reused from the Relations Ontology (RO). | The object and data properties from the ontology are compared to existing RO properties. |
| Documentation (<http://obofoundry.org/principles/fp-008-documented.html>) | The owners of the ontology should strive to provide as much documentation as possible. The documentation should detail the different processes specific to an ontology life cycle and target various audiences (users or developers). | The registry data is checked for ‘homepage’ and ‘description’ entries. If the homepage is present, the URL is checked to see if it resolves (does not return an HTTP status of 400 or greater). |
| Documented Plurality of Users (<http://obofoundry.org/principles/fp-009-users.html>) | The ontology developers should document that the ontology is used by multiple independent people or organizations. | The registry data is checked for ‘tracker’ and ‘usages’ entries. |
| Commitment to Collaboration (<http://obofoundry.org/principles/fp-010-collaboration.html>) | OBO Foundry ontology development, in common with many other standards-oriented scientific activities, should be carried out in a collaborative fashion. | N/A - this cannot be automated at this time. |
| Locus of Authority (<http://obofoundry.org/principles/fp-011-locus-of-authority.html>) | There should be one person responsible for communications between the community and the ontology developers, for communicating with the Foundry on all Foundry-related matters, for mediating discussions involving maintenance in the light of scientific advance, and for ensuring that all user feedback is addressed. | The registry data entry is validated with JSON Schema to ensure that a contact entry is present and that the entry has a name and email address. |
| Naming Conventions (<http://obofoundry.org/principles/fp-012-naming-conventions.html>) | Each entity within the ontology must have a unique label and must not have more than one label. All labels should be declared using the “rdfs:label” property. | ROBOT “report” is run over the ontology. A count of violations for each of the following checks is retrieved from the report: duplicate label, multiple labels, and missing label. |
| Maintenance (<http://obofoundry.org/principles/fp-016-maintenance.html>) | The ontology needs to reflect changes in scientific consensus to remain accurate over time. | A version IRI is retrieved from the ontology and checked against a regex pattern to determine if it is in date format. If so, the date is retrieved to ensure that the ontology is updated in a timely manner. |

**Table 2. OBO Foundry Principles and their automated checks.**
